# Age-specific Jensen contrasts reveal how ontogenetic responses to thermal variability shape lifetime performance in *Daphnia magna*

**DOI:** 10.64898/2026.09.25.754440

**Authors:** Hideyasu Shimadzu, Ilena Day-Dell’Olio, Miguel Barbosa

## Abstract

1. Temperature shapes life histories through its effects on energy allocation among growth, reproduction and maintenance. The resulting temperature dependence of performance is commonly characterised using thermal performance curves (TPCs), which are often treated as static descriptions that apply uniformly throughout life. Yet the response to thermal variability may change across ontogeny.
2. We recorded growth, reproduction and survival throughout the lifetimes of individual *Daphnia magna* under four thermal regimes: Constant Low (15°C), Constant Rearing (20°C), Constant High (25°C) and Variable (15–25°C). The mean temperature under Variable closely matched that under Constant Rearing. Using an allometric energy-allocation model, we derived age-specific and lifetime performance measures and quantified their uncertainty with a stratified individual-level bootstrap. Our principal quantity was the Jensen contrast, defined as performance under Variable minus that under Constant Rearing.
3. Age-specific point estimates of the Jensen contrast changed sign across ontogeny. A positive aggregate contrast in full-trajectory growth concealed a negative mid-life contrast; reproductive point estimates were positive during early and mid-life but negative later. Bootstrap support was strong for the positive early-life and negative mid-life growth contrasts, and for the positive early-life reproductive contrast. Under a second-order approximation, this pattern of estimated signs is consistent with ontogenetic changes in effective thermal curvature.
4. The positive early-life reproductive response did not carry through to the lifetime outcome: expected lifetime reproduction was lower under Variable than under Constant Rearing, with strong bootstrap support for the negative lifetime contrast. Because lifetime reproduction is survival-weighted, lower survival under Variable reduced later reproductive contributions; across the estimated trajectories, the accumulated negative late-life contribution outweighed the earlier positive contributions.
5. These results show why forecasts based on mean temperature, a single life stage or an aggregate TPC may misrepresent lifetime performance under variable thermal regimes. Accounting for both ontogeny and temperature variability may therefore improve forecasts of life-history performance in increasingly variable thermal environments.

## 1 Introduction

Temperature exerts a marked influence on ectotherm life histories through its effects on the allocation of acquired energy among growth, reproduction and maintenance (Savage et al.; 2004; Angilletta Jr.; 2009). The resulting temperature dependence of performance is commonly examined with thermal performance curves (TPCs), which relate a specified measure of organismal performance, such as growth or reproduction, to temperature. A typical TPC is asymmetrically unimodal: performance rises from a critical thermal minimum towards an optimum, after which it declines sharply as temperature approaches a critical maximum (Huey and Stevenson; 1979; Sinclair et al.; 2016). Organisms in nature, however, are seldom exposed to a single constant temperature. As environmental temperature varies, realised performance depends on the portions of the TPC encountered through time rather than on its value at the mean temperature alone. The consequences of thermal variability therefore depend on the shape of the TPC across the temperatures which an organism experiences.

Thermal variability can produce either a positive or a negative departure from performance at the corresponding constant mean temperature (Carrington et al.; 2013; Vasseur et al.; 2014). Jensen’s inequality (Jensen; 1906) relates mean performance under a temperature distribution to performance at its mean; we refer to this difference as the *Jensen contrast*. A positive contrast is consistent with an effectively convex response over the experienced temperature distribution, whereas a negative contrast is consistent with an effectively concave response. Organisms exposed to thermal regimes with identical mean temperatures can therefore differ in average performance when the degree of variability differs. Mean temperature alone cannot determine the sign of the contrast and may provide an incomplete account of performance under a variable thermal regime.

The response to thermal variability, as characterised by the Jensen contrast, need not remain fixed throughout life. Conventional applications often characterise performance at a particular age or life stage and then generalise its thermal dependence across the life course (Sinclair et al.; 2016; Kingsolver and Woods; 2016). Growth, reproduction and lifespan are not, however, independent; ontogenetic shifts in energy allocation bind them together. Many ectotherms broadly conform to the temperature–size rule (Atkinson; 1994): cooler conditions tend to slow growth and prolong life, whereas warmer conditions often accelerate early growth but curtail lifespan (Winkler et al.; 2002; Zuo et al.; 2011). After maturation, energy is increasingly redirected from somatic growth towards reproduction (Angilletta et al.; 2004). Because the thermal sensitivities of these traits reflect both their energetic demands and how energy is allocated among them, the sign of the Jensen contrast may differ between growth and reproduction and may change over the life course. The effective response over the experienced temperature range may therefore be convex at one age yet concave at another. Thermal variability may be associated with higher performance at one age but lower performance at another.

Although the life-stage dependence of thermal responses has long been recognised in principle, few studies have tracked thermal responses across the whole lifespan. Previous work has shown that TPCs can differ among life stages and has emphasised that predictions of ectotherm responses to climate change should account for time-dependent thermal exposure (Sinclair et al.; 2016; Kingsolver and Woods; 2016). Empirical studies likewise show stage dependence: the larval and adult stages of *Littorina obtusata*, for example, differ in both acute and chronic thermal tolerance (Truebano et al.; 2018). It remains uncertain, however, whether the response to thermal variability itself changes sign across ontogeny and how positive and negative age-specific responses combine to determine lifetime performance. If the Jensen contrast changes sign, a snapshot at one age may indicate the opposite response at another and need not predict the lifetime outcome. The consequences of thermal variability can therefore be understood at three connected levels. First, fluctuations expose organisms to different portions of a nonlinear temperature-response function, represented here by a TPC, and thereby generate a Jensen contrast. Second, the effective thermal response may change across ontogeny as energy allocation among growth, reproduction and maintenance shifts. Third, lifetime performance integrates positive and negative age-specific contributions, so that the sign observed locally need not predict the lifetime outcome.

We examine these connected levels in *Daphnia magna*, a model ectotherm whose growth, reproduction and lifespan are strongly temperature dependent. We followed individuals from birth to death under Constant Low, Constant Rearing, Constant High and Variable temperature treatments. The mean temperature under Variable closely matched that under Constant Rearing; the three constant-temperature treatments provide reference observations for the direction of temperature dependence. Our principal comparison is between Variable and Constant Rearing, and the sign of the resulting contrast indicates whether performance under thermal variability is higher or lower than under a constant regime with a nearly identical mean temperature.

Here, we ask (1) how the sign and magnitude of Jensen contrasts in growth vary with age; (2) whether the corresponding contrasts in survival-weighted reproduction follow the same ontogenetic pattern; and (3) whether full-trajectory growth and lifetime reproduction preserve or conceal the age-specific contrasts from which they arise. To address these questions, we use the allometric energy-allocation model (Shimadzu and Barbosa; 2025) to derive age-specific and lifetime performance quantities from the treatment-specific trajectories and quantify the corresponding Jensen contrasts. We examine survival separately to clarify its contribution to lifetime reproduction. Together, these analyses distinguish age-specific responses to thermal variability from the lifetime outcome that emerges as those responses accumulate. We show that the estimated age-specific Jensen contrasts can reverse sign across ontogeny and that a positive early-life reproductive response does not predict the sign of the lifetime reproductive outcome.

## 2 Materials and methods

The experimental data and the allometric energy-allocation model have been described previously (Shimadzu and Barbosa; 2025). Here, we summarise the experimental procedures and define the theoretical performance quantities required for the present Jensen-contrast analysis.

### Study organism and parental generation

All experimental individuals were F_1_ descendants of fourth-brood neonates from *D. magna* clone F (Baird et al.; 1991), which has a documented sensitivity to environmental stress (Barbosa et al.; 2014, 2015). The parental generation (F_0_; *N* = 30) was maintained at 20°C under a 16:8-hour light:dark photoperiod in ASTM medium and fed the green alga *Pseudokirchneriella subcapitata* every second day at 3.0 *×* 10^5^ cells ml^*−*1^; the medium was renewed on the same schedule. Culture conditions throughout complied with the OECD guidelines for reproduction tests with *Daphnia* (OECD; 2012).

### Temperature treatments and climate change rationale

Immediately after birth, F_1_ individuals were placed in individual 50-ml glass containers and randomly assigned to one of four lifetime temperature treatments in a Binder BS28 incubator: Constant Low (15°C; *N* = 158), Constant Rearing (20°C; *N* = 156), Constant High (25°C; *N* = 157) and Variable (15–25°C; *N* = 157). Constant Rearing represents standard laboratory maintenance conditions. The bounds of 15°C and 25°C bracket the reported thermal optimum of approximately 20–21°C, yet remain below the temperatures at which mortality rises sharply (Giebelhausen and Lampert; 2001; Maceda-Veiga et al.; 2015).

The Variable treatment was devised to reflect diurnal thermal dynamics rather than an arbitrary fluctuation. Within each day, temperature varied stochastically between 15°C and 20°C during the dawn–morning and late-afternoon periods (00:00–08:00 and 18:00–24:00), and between 20°C and 25°C during the morning–afternoon period (08:00–18:00). The overall mean under Variable was 19.8°C, nearly identical to that under Constant Rearing (20°C); for the Jensen-contrast analysis, we regarded both means as 20°C. The comparison between these treatments therefore primarily reflects temperature variability and its attendant thermal history rather than a difference in mean thermal exposure.

### Life-history measurements

All F_1_ individuals received the feeding regime described for F_0_, and the medium was renewed every second day throughout the experiment. For each individual, we recorded (1) body length at birth (*t* = 0), at every subsequent brood event and at death (*t* = *τ*); (2) the number of F_2_ neonates produced at each brood event; and (3) the interval between successive brood events. Because body length was recorded at brood events, observations were irregularly spaced in time rather than occurring at fixed intervals.

At each measurement, we transferred each individual to a culture plate with a 3-ml plastic pipette and photographed it. We used ImageJ (Schneider et al.; 2012) to measure body length in millimetres from the tip of the head to the base of the caudal spine. The experiment continued until the last surviving F_1_ individual died on day 141. Altogether, we recorded more than 130,000 F_2_ neonates across the four treatments.

### Replication statement

Each temperature treatment was maintained in a separate, identical incubator (Binder BS28) throughout the experiment, and glass containers were not rotated between incubators. Temperature was applied at the incubator level, with one incubator per treatment, and individuals within an incubator are subsamples rather than independent replicates of the thermal regime (Table 1). To minimise confounding between incubator and treatment, all other conditions (medium, food ration, renewal schedule, photoperiod and handling) were identical across incubators. F_1_ individuals were produced by a common pool of 30 F_0_ females and randomly allocated to their temperature treatments, so that maternal effects were distributed across treatments rather than aligned with them. Incubator temperatures were monitored with temperature loggers throughout. Our inferences concern individual-level responses to the four thermal regimes. The individual-level bootstrap quantifies uncertainty among individuals but cannot separate treatment effects from incubator effects.

**Table 1:** Replication statement for the temperature experiment.

| Scale of inference | Scale at which the factor of interest is applied | Number of replicates at the appropriate scale |
| --- | --- | --- |
| Individual | Incubator (one per temperature treatment) | 1 incubator per treatment; within each: Constant Low, $N = 158$; Constant Rearing, $N = 156$; Constant High, $N = 157$; Variable, $N = 157$ individuals |

### Model structure and theoretical performance quantities

Our performance measures derive from an allometric model in the tradition of the Bertalanffy–Pütter growth equation (Pütter; 1920; von Bertalanffy; 1934, 1957), in which growth represents the balance between anabolic production and maintenance expenditure. Subsequent extensions partition net production between somatic growth and reproductive investment (Quince et al.; 2008; Mollet et al.; 2010; Shimadzu and Wang; 2022) and express the resulting trajectories in terms of body length (Shimadzu and Barbosa; 2025).

We use this theoretical structure to define the age-specific, full-trajectory and lifetime performance quantities from which we quantify Jensen contrasts.

For each temperature treatment *c*, the model describes body-length growth as

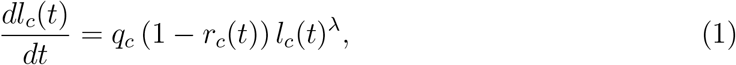

where *q*_*c*_ is a treatment-specific coefficient that governs net production, *λ* is the allometric exponent shared among treatments, and *r*_*c*_(*t*) is the time-varying relative allocation to direct reproduction (Shimadzu and Barbosa; 2025).

### Growth

The instantaneous somatic growth rate is

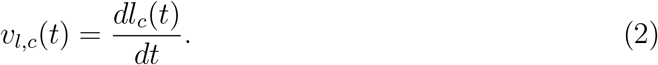

We also define a descriptive full-trajectory growth rate from birth to the last age supported by the trajectory in each treatment. When compared across treatments, this aggregate quantity yields a conventional TPC-like summary, but the treatment-specific endpoints preclude comparison over a common age interval. We therefore use it as a descriptive biological summary and a foil for the age-specific analysis, rather than as an estimate of age-invariant thermal performance.

### Reproduction and survival

The model connects body-length growth to reproductive investment through *r*_*c*_(*t*) (Equation 1). The corresponding instantaneous model-derived reproductive-energy rate is

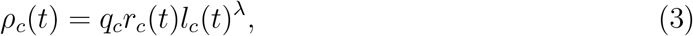

and its integration over time gives cumulative reproductive energy, 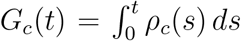. We relate this latent quantity to the expected cumulative number of neonates up to time *t* through

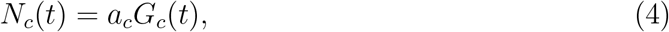

where *a*_*c*_ is a treatment-specific coefficient that converts reproductive energy into neonate number. Letting *τ*_*c*_ denote individual lifespan, we write the treatment-specific survival function as *S*_*c*_(*t*) = Pr(*τ*_*c*_ *> t*). The rate of neonate production is then

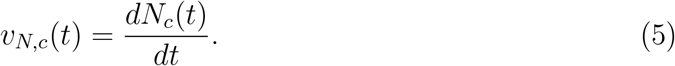

We combine this conditional production rate with survival to define the survival-weighted reproductive rate, which represents the age-specific contribution to expected lifetime reproduction:

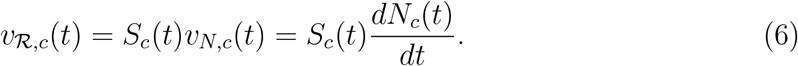

Expected lifetime reproduction (Dublin and Lotka; 1925), equivalently the net reproductive rate, is then

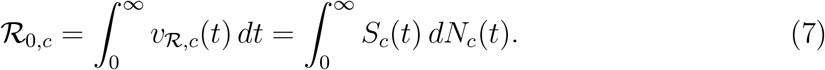

To clarify the survival component of expected lifetime reproduction, we also summarise each treatment by its mean lifespan, 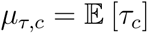.

### Jensen contrasts

For a generic performance rate *v*_*P*_ (*t*), where *P* denotes growth or reproduction, we define the theoretical age-specific Jensen contrast as the difference between Variable and Constant Rearing:

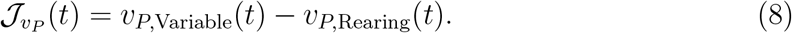

The lifetime reproduction contrast can equivalently be expressed as the signed accumulation of the age-specific rate contrasts:

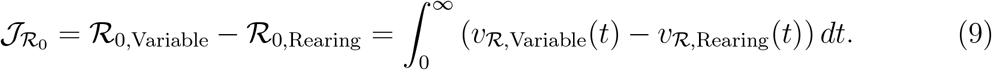

The contrast within any one age window therefore need not share the sign of the lifetime contrast, which depends on the signs, magnitudes and durations of contributions across the complete lifespan.

Under Constant Rearing, temperature is fixed at *µ* = 20°C, whereas under Variable, temperature *C* is a random variable with a mean of 19.8°C, which we approximate as 20°C for the present analysis. Equation (8) therefore describes the effect of temperature variability at an approximately matched mean. To place this contrast within the thermal-performance framework, let *f*_*t*_(*C*) denote the effective TPC at age *t*. A second-order Taylor expansion around *µ* = 20 gives the following approximation to expected performance under Variable:

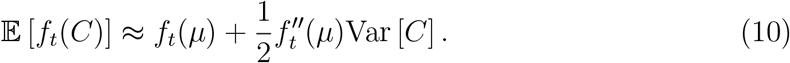

It follows from Equation (10) that the Jensen contrast is

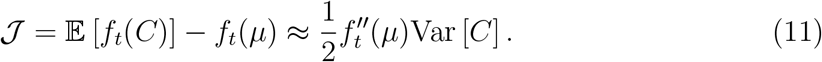

Under this second-order approximation, the age-specific Jensen contrast has the same sign as the second derivative of the local TPC, 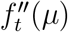. A positive contrast is therefore consistent with 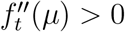 and with a response that is effectively convex at age *t*, whereas a negative contrast is consistent with 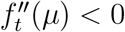 and with one that is effectively concave. Under the same approximation, a reversal in the sign of the contrast across age is likewise consistent with an ontogenetic change in effective thermal curvature.

We use Equation (11) as an interpretive approximation for the age-window growth and reproductive contrasts, rather than as an exact estimator of instantaneous TPC curvature. The range from 15 to 25°C may not be sufficiently narrow for higher-order terms to be negligible; the temperature distribution may be asymmetric; and the observed contrast may incorporate the effects of thermal history, acclimation, survival and energy allocation. We therefore regard positive and negative age-window contrasts as evidence of *effective* convexity and concavity, respectively. Full-trajectory growth and lifetime reproduction are integrative empirical comparisons, and we do not interpret their contrasts as estimates of curvature at any single age. The sign and magnitude of the Variable-minus-Constant Rearing contrast, rather than a curve fitted through three constant-temperature points, provide the primary evidence for the direction of the response to thermal variability.

### Estimation and uncertainty

We fitted Equation (1) by gradient matching and reconstructed the treatment-specific growth and reproductive trajectories from which we derived the performance summaries and Jensen contrasts reported here. Hats denote estimated quantities. Appendix A presents the fitted allocation trajectories.

Treatment-specific survival functions were estimated with the Kaplan–Meier estimator (Kaplan and Meier; 1958); no observations were censored. We quantified uncertainty with a nonparametric bootstrap stratified by temperature treatment, in which complete individual records were resampled with replacement and the model, survival functions and all derived quantities were re-estimated in each of 1,000 replicates. We report 95% bias-corrected (BC) bootstrap confidence intervals together with the empirical proportions of replicates in which each Jensen contrast was positive or negative; these proportions are not posterior probabilities.

## 3 Results

### 3.1 Age-specific Jensen contrasts in growth

The body-length trajectories were broadly similar across treatments, although the trajectory under Constant Low was distinct from the others (Figure 1A). Daily growth increments declined rapidly during early life, and the fitted trajectories crossed at approximately days 12 and 42 (Figure 1B). We therefore used these crossings to define three data-derived descriptive age windows: days 0–12, 12–42 and 42 onwards. These windows do not necessarily correspond to independently defined developmental stages; for growth, the final window ended at day 79, the last age for which fitted trajectories were supported in all treatments.

**Figure 1:**
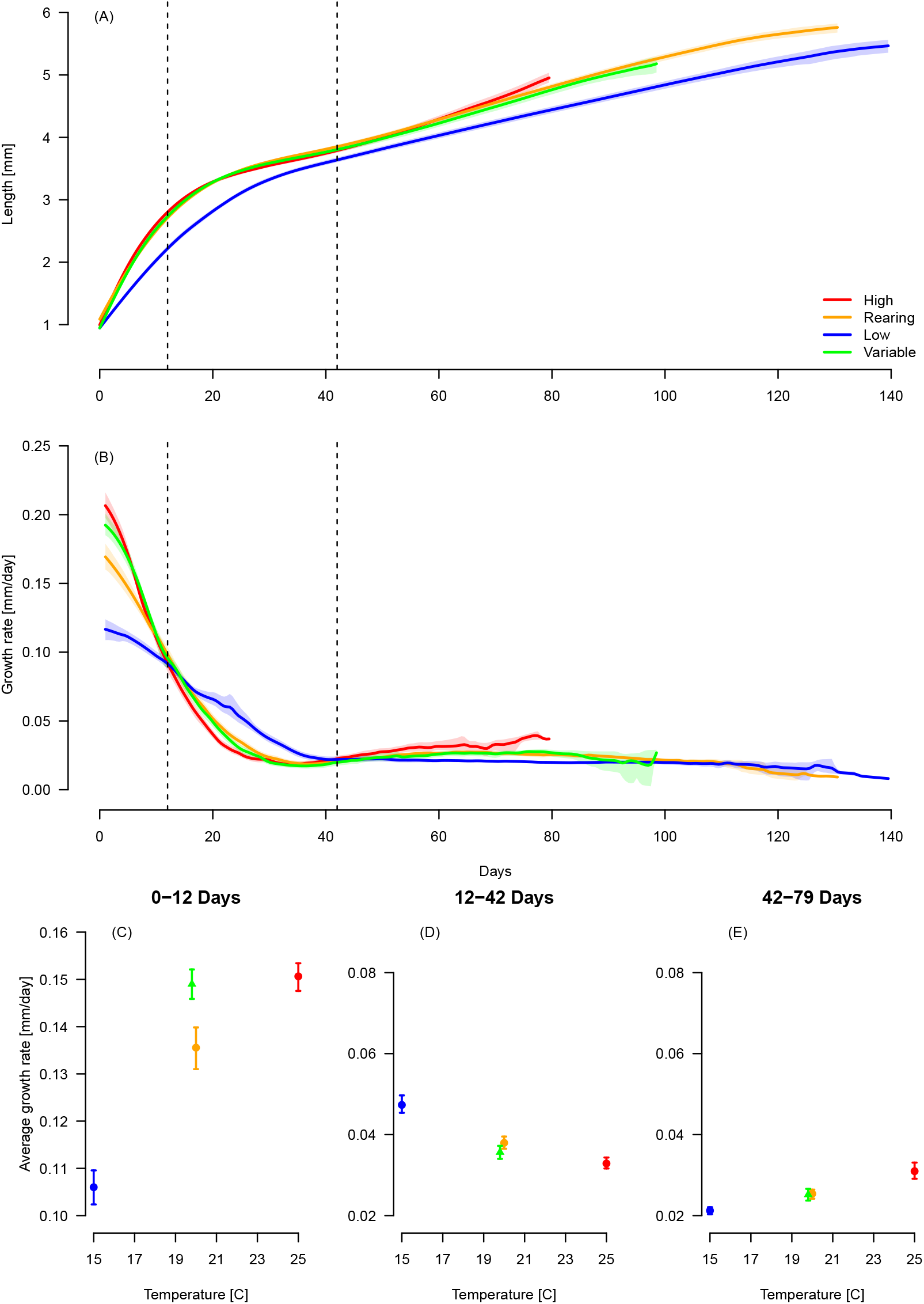
(A) Body-length growth trajectories under each temperature treatment. (B) Daily growth increments under each treatment. Dashed vertical lines mark *t* = 12 and *t* = 42, at which the fitted growth-rate trajectories cross. (C–E) Mean growth rates over the age windows spanning days 0–12, 12–42 and 42–79, respectively. The crossings in Panel B define these data-derived descriptive windows. In Panels C–E, Constant Low, Constant Rearing and Constant High provide the constant-temperature reference values. The Variable estimate represents the response integrated over the 15–25°C temperature distribution; its difference from Constant Rearing gives the Jensen contrast. Shading and error bars denote pointwise and scalar 95% bias-corrected bootstrap intervals, re-spectively.

During the 0–12-day window, growth was slower at lower constant temperatures and faster at higher ones (Figure 1C). This ordering reversed during days 12–42 and returned to its initial direction during days 42–79 (Figure 1D–E). The pattern suggests that the direction of temperature dependence changed with age, although the three constant-temperature observations alone could not identify the curvature of a continuous TPC.

The growth Jensen contrast was 0.0135 mm day^*−*1^ during days 0–12 (95% BC interval: 0.0079, 0.0187; all bootstrap contrasts positive), but *−*0.00230 mm day^*−*1^ during days 12–42 (95% BC interval: *−*0.00464, *−*0.00006; 98.0% of bootstrap contrasts negative). During days 42–79, the contrast was small and uncertain (*−*0.00022 mm day^*−*1^; 95% BC interval: *−*0.00212, 0.00154; Figure 1C–E). The estimated growth contrast therefore changed direction across ontogeny. Under the second-order approximation in Equation (11), these well-supported signs are consistent with effective convexity during the first age window and effective concavity during the second.

### 3.2 Age-specific Jensen contrasts in reproduction

Treatment differences were more pronounced in the fitted cumulative numbers of neonates (Figure 2A) than in the body-length trajectories. The trajectories under Constant High and Variable were similar. Survival-weighted reproductive rates rose towards treatment-specific peaks and declined as survival diminished (Figures 2B and 4); their relative positions changed at approximately day 42.

**Figure 2:**
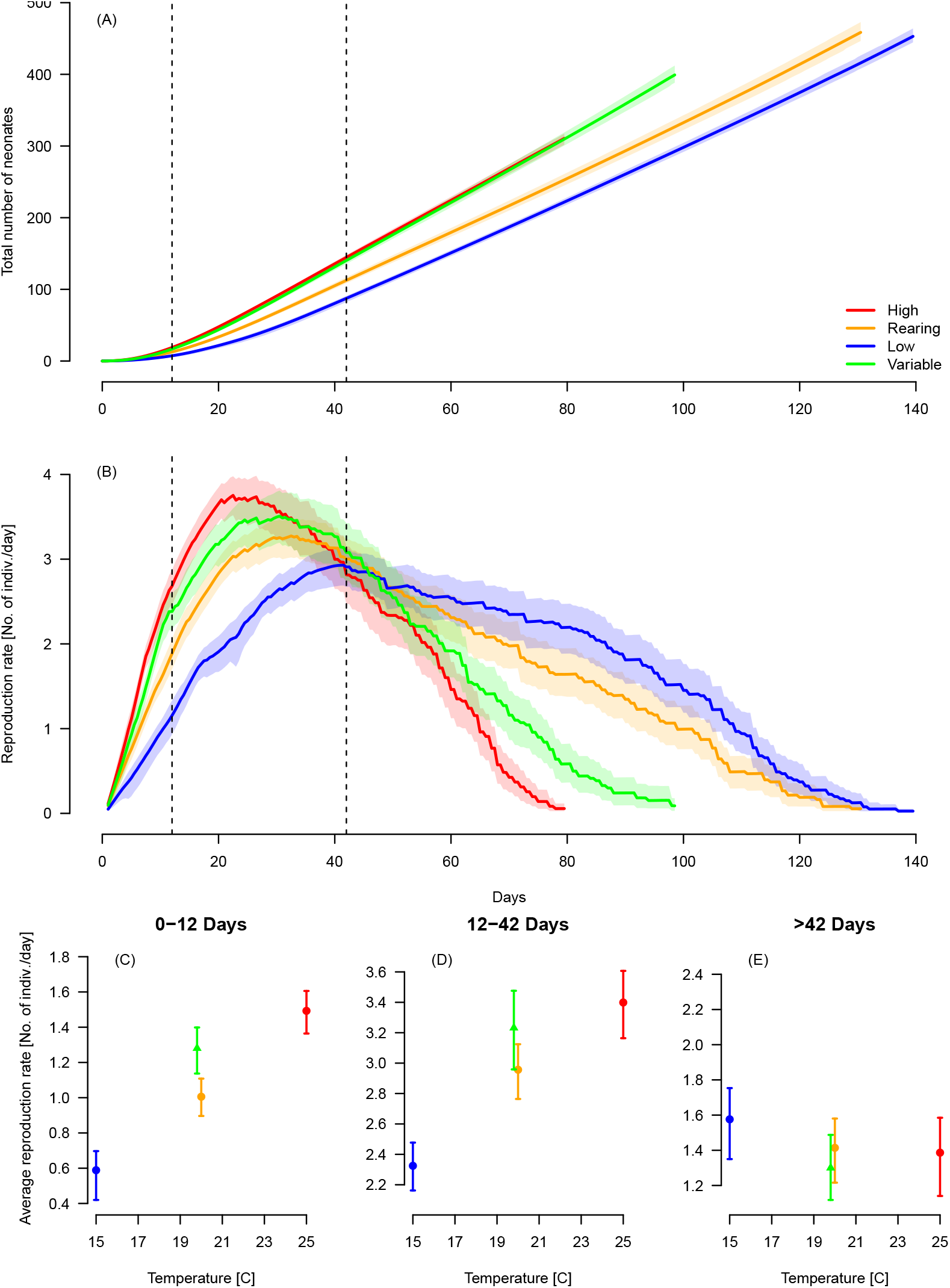
(A) Model-derived expected cumulative numbers of neonates under each temperature treatment. (B) Daily survival-weighted reproductive rates (Equation 6) under each treatment. Dashed vertical lines mark *t* = 12 and *t* = 42, at which the fitted growth-rate trajectories cross (Figure 1B). (C–E) Mean survival-weighted reproductive rates over the age windows spanning days 0–12, 12–42 and all supported ages after day 42, respectively. The growth-rate crossings in Figure 1B define these data-derived descriptive windows. In Panels C–E, the difference between Variable and Constant Rearing gives the Jensen contrast. Shading and error bars denote pointwise and scalar 95% bias-corrected bootstrap intervals, respectively.

At constant temperatures, average survival-weighted reproductive rates were higher under warmer conditions during days 0–12 and 12–42 (Figure 2C–D). The Variable-minus-Constant Rearing contrast was 0.275 neonates day^*−*1^ during days 0–12 (95% BC interval: 0.117, 0.413; all bootstrap contrasts positive) and 0.275 neonates day^*−*1^ during days 12– 42 (95% BC interval: *−*0.044, 0.572; 95.5% of bootstrap contrasts positive). Beyond day 42, the contrast reversed sign, with a point estimate of *−*0.114 neonates day^*−*1^ (95% BC interval: *−*0.376, 0.164; 82.8% of bootstrap contrasts negative; Figure 2E). These point estimates changed sign across age, although bootstrap support for the middle and late contrasts was weaker than that for early reproduction.

### 3.3 Full-trajectory and lifetime integration

When growth was summarised across each treatment’s full fitted trajectory, the three constant-temperature observations formed a familiar aggregate TPC-like response in which average growth was greater at warmer temperatures (Figure 3A). Growth under Variable exceeded that under Constant Rearing by 0.00714 mm day^*−*1^ (95% BC interval: 0.00532, 0.00888; all bootstrap contrasts positive). This whole-trajectory result resembled the early-life contrast, yet concealed the negative contrast during days 12–42 and the near-zero contrast during days 42–79. Because the fitted endpoints differed among treatments, this whole-trajectory quantity is a descriptive summary of each supported trajectory rather than a common-age estimate.

**Figure 3:**
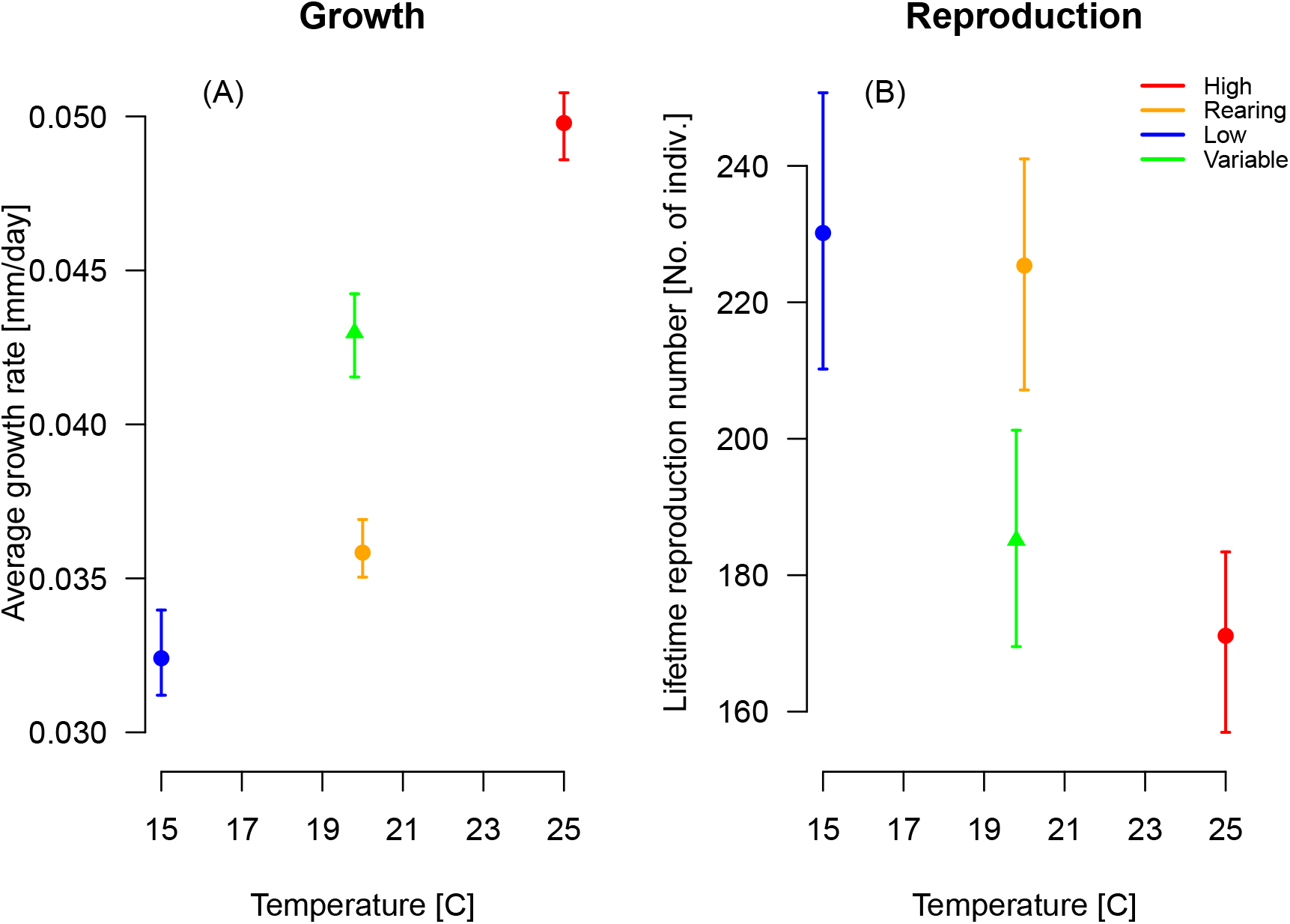
Aggregate thermal-performance summaries for (A) growth across the full fitted trajectory and (B) expected lifetime reproduction, 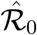. The three constant-temperature values provide reference points, whereas the difference between Variable and Constant Rearing gives the Jensen contrast. Panel A shows the TPC-like summary obtained when growth is aggregated across ontogeny. Because the fitted trajectories end at 79.5, 98.5, 130.5 and 139.5 days for Constant High, Variable, Constant Rearing and Constant Low, respectively, Panel A is a descriptive whole-trajectory summary rather than a common-age comparison. Error bars denote 95% bias-corrected bootstrap intervals.

In contrast to aggregate growth, lifetime reproduction yielded a negative integrated contrast (Figure 3B). Expected lifetime reproduction was 185.1 neonates under Variable and 225.4 under Constant Rearing, giving a Jensen contrast of *−*40.3 neonates (95% BC interval: *−*64.3, *−*17.4; 99.9% of bootstrap contrasts negative). As Equation (9) formalises, this contrast is the signed accumulation of the age-specific contributions. Lower survival under Variable reduced later reproductive contributions; the accumulated negative contribution after day 42 outweighed the earlier positive contributions. An early-life snapshot would therefore have suggested a lifetime outcome opposite in sign to that estimated over the full life history.

Survival differed markedly between Variable and Constant Rearing (Figure 4). Mean lifespan was 51.9 days under Variable (95% BC interval: 48.1, 56.1) and 72.0 days under Constant Rearing (95% BC interval: 66.8, 76.9); the Variable-minus-Constant Rearing difference was *−*20.0 days (95% BC interval: *−*26.5, *−*13.6; all bootstrap differences negative). Constant High had a similarly short mean lifespan of 48.4 days, whereas Constant Low had the longest, at 80.9 days.

**Figure 4:**
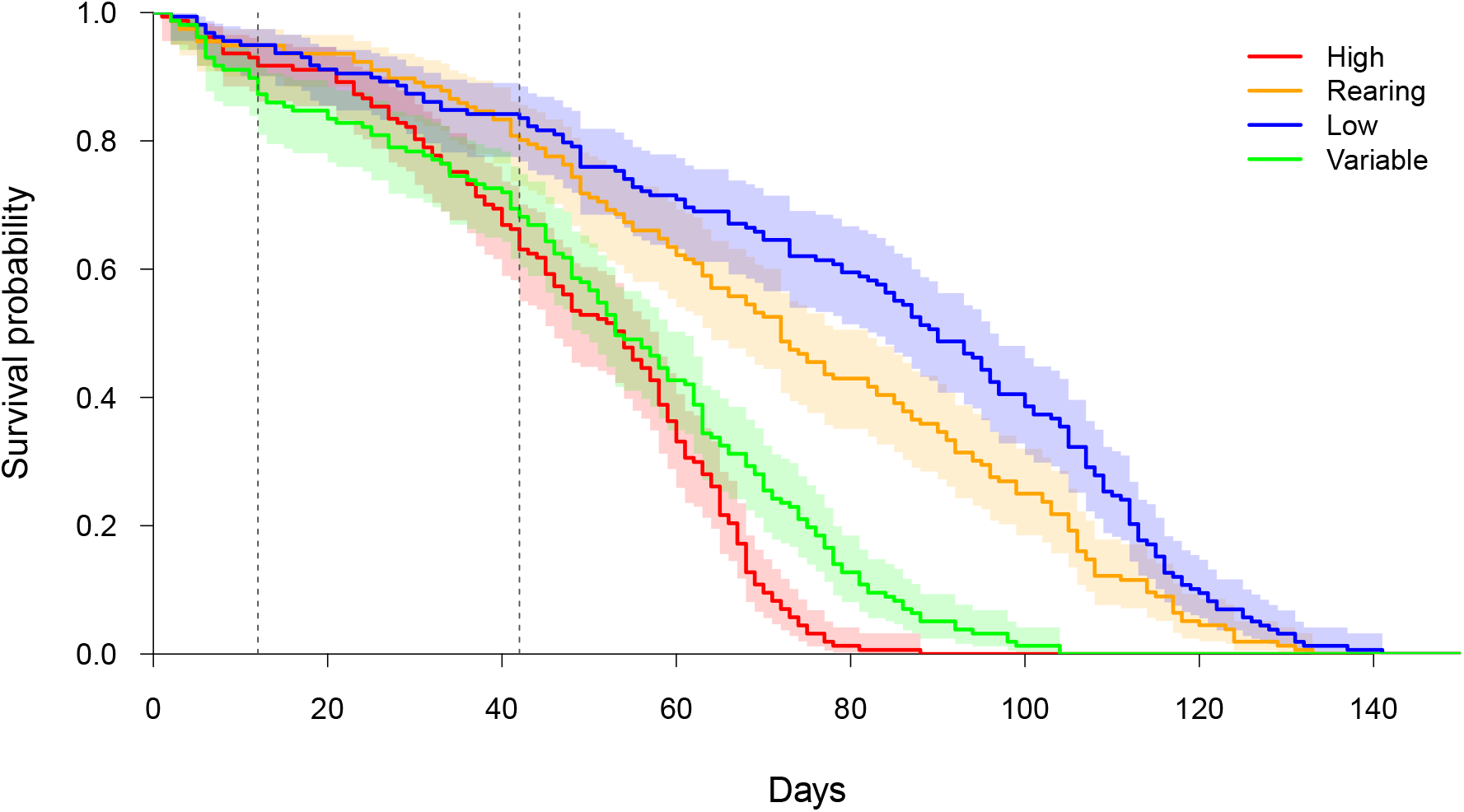
Kaplan–Meier survival curves for the four temperature treatments. All deaths were observed; shaded regions denote pointwise 95% confidence intervals. Dashed vertical lines mark days 12 and 42, which delimit the descriptive age windows used for the growth and reproductive analyses.

## 4 Discussion and conclusion

The aggregate and age-specific growth analyses yielded different but complementary accounts. When growth was summarised over each treatment’s full estimated trajectory, average growth increased from Constant Low to Constant High, and Variable exceeded Constant Rearing, producing a familiar aggregate TPC-like response. Yet this aggregate comprised an early positive Jensen contrast, a mid-life negative contrast and a small late contrast; the growth Jensen contrast was therefore not age invariant. Although the full trajectories ended at treatment-specific ages and did not permit a strict common-age comparison, they remain informative because they represent the aggregate view that can obscure ontogenetic reversals. The age-window analysis revealed those reversals.

Age-specific reproductive contrasts likewise did not necessarily predict the lifetime outcome. Variable was associated with higher reproductive performance during early life, and the mid-life contrast also had a positive point estimate; yet expected lifetime reproduction was approximately 40 neonates lower than under Constant Rearing. This apparent discord is not a contradiction. Equation (9) shows that the lifetime contrast is the signed accumulation of age-specific reproductive contributions, each weighted by survival. Survival declined more rapidly under Variable, reducing the contribution of later reproduction; mean lifespan was approximately 20 days shorter than under Constant Rearing. Across the fitted trajectories, the accumulated negative contribution after day 42 outweighed the earlier positive contributions. An early-life snapshot would therefore have suggested a lifetime outcome opposite in sign to that estimated over the full life history.

Our central inference concerns age-dependent changes in the response to temperature variability, rather than the curvature of a TPC inferred from the three constant-temperature observations alone. Those observations show whether performance increases or decreases across 15, 20 and 25°C, but cannot robustly identify curvature. The Variable treatment provides a different comparison because its mean nearly matches that of Constant Rearing. When the higher-order remainder is modest, Equation (11) relates the Variable–Constant Rearing difference to temperature variance and the local second derivative. Because the experimental distribution spans 15–25°C and biological responses may retain thermal history, we interpret the signs of the age-window contrasts as evidence of effective convexity or concavity over the experienced distribution, not as proof of pointwise instantaneous curvature. We do not extend this local-curvature interpretation to the integrative full-trajectory and lifetime-reproduction contrasts.

The allocation model (Equation 1) provides a conceptual link between growth and reproduction but is not required for the Jensen argument itself. Within the fitted model, the estimated allocation trajectories, 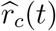, generally increased with age and approached a plateau; this pattern reflects a relative shift from somatic growth towards reproduction (Figure A1). Because these trajectories are inferred from the same body-length records used to estimate growth, they provide model-based context rather than independent evidence for the mechanism. The allocation model was originally developed to characterise differences in life-history trajectories among thermal regimes (Shimadzu and Barbosa; 2025); here we use the same structure to ask how variability at an approximately matched mean changes performance with age and how those age-specific changes accumulate into lifetime outcomes.

Thermal history, acclimation and repeated exposure to the upper portion of the performance range may likewise contribute to the age dependence of the Jensen contrasts. Higher temperature commonly increases metabolic demand and promotes earlier investment in growth and reproduction at the expense of lifespan (Winkler et al.; 2002; Schulte; 2015), whereas short-term fluctuations can alter growth and resource constraints in *Daphnia* (Balseiro et al.; 2021). The reported thermal optimum for *D. magna* is approximately 20–21°C (Giebelhausen and Lampert; 2001), and mortality rises sharply above 26°C (Maceda-Veiga et al.; 2015). Under Variable, individuals experienced 20–25°C for ten hours each day and therefore repeatedly encountered temperatures at or above the reported thermal optimum. The Variable treatment was structured around diurnal thermal dynamics rather than an arbitrary fluctuation; the observed contrasts should therefore be interpreted in the context of repeated thermal exposure and thermal history, rather than attributed to temperature variance alone.

The resemblance between Variable and Constant High was consistent with “cryptic warming” pattern, in which life-history activity shifts towards earlier ages at the expense of longevity and lifetime reproduction. In descriptive terms, *D. magna* under Variable exhibited a “jack of all temperatures, master of none” pattern: performance was broadly comparable with that under the constant regimes without consistently exceeding them, while both longevity and lifetime reproduction were lower than under Constant Rearing. This pattern may also relate to bet-hedging under unpredictable environments. Bet-hedging theory predicts that selection may favour reduced variance in fitness across generations, even at the expense of arithmetic mean fitness (Seger and Brockmann; 1987; Simons; 2011), whereas unpredictable temperature has previously been associated with greater variance in reproductive investment in *D. magna* (Barbosa et al.; 2015). The present experiment did not, however, test whether the observed contrasts represented an adaptive response or a physiological consequence of thermal history; multigenerational experiments would be required to distinguish these possibilities.

Age-specific Jensen contrasts connect two questions that aggregate TPCs treat separately: how a temperature distribution changes performance at a given age, and how such local changes are integrated across the life history. The combination of longitudinal growth, reproduction and survival records permits both questions to be addressed without conflating them. In this experiment, early growth and reproduction were higher under Variable than under Constant Rearing, whereas lifetime reproduction and lifespan were lower. Whether the variable regime appeared beneficial or costly therefore depended on when performance was measured and on how the resulting contributions accumulated. Together, these results suggest that the consequences of thermal variability are best understood at three connected levels: the age-specific Jensen contrast, its change across ontogeny and the accumulation of age-specific contributions into lifetime performance. A Jensen framework that accounts for all three levels may improve forecasts of life-history performance under variable thermal regimes.

## Appendix A: Graphical presentation of 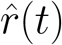

**Figure A1:**
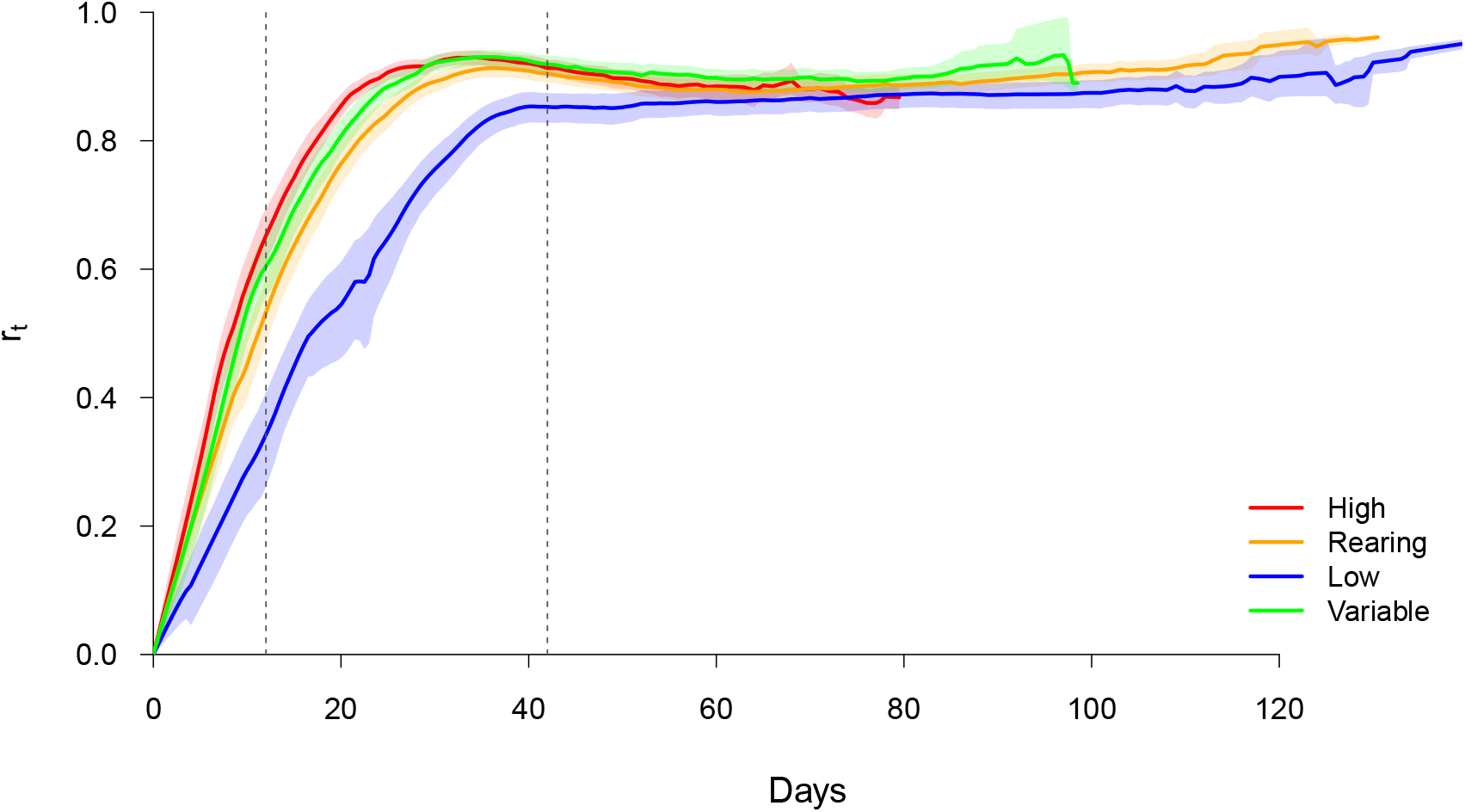
Model-derived relative allocation to direct reproduction, 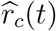, under the four temperature treatments. The fitted trajectories generally increase with age and approach a plateau. Shaded regions denote pointwise 95% bias-corrected bootstrap intervals; dashed vertical lines mark days 12 and 42. These trajectories describe an internal model component that links growth and reproduction; they do not provide an independent test of the mechanism underlying the Jensen contrasts.

